# Diverse Secondary Metabolites in Methanolic and Alkaloidal Extracts of *Hunteria umbellata* Leaves: Insights from Computational Molecular Networking

**DOI:** 10.1101/2024.09.18.613617

**Authors:** Rukayat A. Adedeji, Musibau Opemipo, Oluwabukunmi Babalola, Blessing Titilayo, Solomon O. Julius, Stephenie C. Alaribe

**Affiliations:** Department of Pharmaceutical Chemistry, Faculty of Pharmacy, University of Lagos, Nigeria; Department of Pharmacognosy, Faculty of Pharmacy, University of Lagos, Nigeria

**Keywords:** Medicinal Plant, *Hunteria umbellata*, Computational Molecular Networking, LC-MS/MS, Omics

## Abstract

Plants have long served as a vital source of therapeutic agents in both traditional and orthodox medicine. However, with the shift in drug discovery towards laboratory synthesis, there is a decline in the exploration of natural sources for drug development. This downturn calls for return to natural drug discovery and, more importantly, towards the development of improved methods of isolating, identifying, and characterising chemical moieties obtained from plants. This study redirects attention to natural product research by employing advanced metabolomic and computational approaches to characterise the bioactive compounds of *Hunteria umbellata*. Metabolomic technique was employed, utilising liquid chromatography-tandem mass spectrometry (LC-MS/MS) to separate the chemical components. The isolated compounds were then identified using their mass-to-charge (m/z) ratios. Chromatograms were analysed using a computational molecular networking tool to match the m/z values to known compounds in mass spectrometry libraries. Eighteen compounds were successfully isolated from the methanolic and alkaloidal extracts, including Yohimbine, (−)-Epicatechin, Picrinine, Tubotaiwine, Quercetin-3-O-robinobioside, and Pheophorbide A. To our knowledge, this represents the first comprehensive metabolomic profiling of *H. umbellata* using computational molecular networking, revealing a diverse set of flavonoids and indole alkaloids. Notably, the detection of Pheophorbide A, a chlorin derivative with photodynamic therapy potential, constitutes a new report for this species and suggests unexplored therapeutic relevance. These findings provide significant insight into the bioactive components of *Hunteria umbellata*, supporting its traditional medicinal uses. Identifying clinically relevant compounds not only validates traditional practices but also highlights the plant’s potential for contributing to modern drug discovery efforts.

## INTRODUCTION

Plants are an important source of medicine and play a key role in world health (Aslam and Ahmad, 2016; Chopra and Dhingra, 2021). Medicinal plant use is ever on the rise as they have been known to be an important source of everyday medicine and groundbreaking discoveries in the treatment of cancer, Alzheimer’s, sickle-cell anaemia, and certain autoimmune diseases. Plants remain an important and continuing source of novel chemotherapies even though the majority of modern drug discoveries are a result of synthetic compounds and monoclonal antibodies development (Ouyang *et al*., 2014; Dehelean *et al*., 2021).

*Hunteria umbellata* K. Schum, a tropical rainforest tree found in western and central Africa, is a member of the Apocynaceae family (Adeneye and Adeyemi, 2009; Igbe *et al*., 2009a; Oboh *et al*., 2019). The Apocynaceae family is one of the most extensively studied plant families, and many genera, including Hunteria, have been studied for their chemical composition, pharmacological activity and economic importance. Members of the Hunteria genus are quite popular amongst traditional practitioners, and many of their uses in herbal medicine preparation have been justified in scientific studies. For instance, a study on the anti-inflammatory activity of *Hunteria zeylanica* found that crude alkaloid extracts of the plant’s stem bark significantly inhibited acute inflammation (Igbe *et al*., 2009b).

The major challenge with incorporating herbal products into the national healthcare plan in African countries is the lack of unbiased, concrete scientific evidence to ascertain their identity, purity, and quality (Falodun 2010). The majority of the current analytical techniques available for identifying components of herbal products are not sensitive to the complex nature of these products. Thus, the inadequacy in properly accounting for all the chemical constituents present in a plant preparation is one of the major reasons why people do not trust its efficacy or safety.

To fully elucidate the chemical complex of a natural plant/product, it is important to use an analytical technique that goes beyond the traditional methods of revealing the structural components. Various omics technologies have revolutionised how drugs of biological origin are discovered and processed to produce efficacious and high-quality medicines. Metabolomics tools like liquid chromatography-tandem mass spectrometry (LC-MS/MS) use mass data of components to exhaustively elucidate these components beyond what simple nuclear magnetic resonance (NMR) would do. LC-MS/MS data will account for the total ions present and thus would identify any compound or fragments attached to a major compound.

Global Natural Products Social Molecular Networking (GNPS; http://gnps.ucsd.edu) is an open-access knowledge base for the community-wide organisation and sharing of raw, processed, or identified tandem mass (MS/MS) spectrometry data. This study uses this platform as a computational molecular networking tool for analysing the spectra from LC-MS/MS analysis. GNPS is a data-driven platform for the storage, analysis, and dissemination of MS/MS spectra knowledge (Wang *et al*., 2016). It enables the community to share raw spectra, continuously annotate deposited data, and collaboratively curate reference spectra (referred to as spectral libraries) and experimental data (organized as datasets) (Wang *et al*., 2016). GNPS provides the ability to analyse a dataset and compare it to all publicly available data. Molecular networks are visual representations of the chemical space present in MS experiments. GNPS is employed for molecular networking, a spectral correlation and visualisation approach that detects spectra from related molecules (so-called spectral networks), even when the spectra are not matched to any known compounds. Spectral alignment detects similar spectra from structurally related molecules, assuming these molecules fragment in similar ways, as reflected in their MS/MS patterns, analogous to the detection of related protein or nucleotide sequences by sequence alignment (Wang *et al*., 2016). This tool enables a cycle of annotation in which users curate data, enabling product identification, and a knowledge base of reference spectral libraries and public data sets is created.

The objective of this study is to identify and characterise the secondary metabolites present in the methanolic and alkaloidal extracts of *Hunteria umbellata* using LC-MS/MS and computational molecular networking tool.

## MATERIALS AND METHODS

### Materials

Fresh leaves of *Hunteria umbellata* were collected for this study. The chemicals used for extraction included 98% methanol, 100% chloroform, 2M sulfuric acid, and ammonia. Experimental procedures were carried out using the following laboratory equipment: a LC-MS/MS setup, a BÜCHI Rotavapor® rotary evaporator, a Swallow Laboratory Oven (LTE Scientific Ltd., United Kingdom), a Christy Laboratory Hammer Mill Grinder (Christy Turner Ltd., England), and an OHAUS Scout Balance, Model SPX621. Additional materials included glass funnels, beakers, foil paper, stirring rods, Whatman No. 1 filter paper (32.0 cm, CAT No: 1001320, Whatman England), spatulas, and ziploc bags.

### Collection and Identification

Fresh samples of the leaves and stem bark of *Hunteria umbellata* were collected and identified by a taxonomist in Ibadan, Oyo State, Nigeria. The sample was transported to Lagos, identified and authenticated by Dr Nodza George at the herbarium in the University of Lagos, Akoka. Voucher number LUH-8748 was obtained.

### Plant Material Preparation

The leaves were rinsed thoroughly using distilled water to remove sand, dust, and any adhering particles from the leaf surface, after which the leaves were flapped to drain most of the water and then spread out on a flat, clean surface inside the lab, near the window. The leaves were then plucked from the stem and spread out to air dry at room temperature for 8 days, during which the leaves were regularly turned over for optimum drying. The dried leaves were placed in the hot air oven for 30 minutes before grinding to make the leaves material crunchy enough for the milling process. Afterwards, the crispy leaves were crushed by hand to increase the surface area of the materials. A lab mill grinder was used to grind the leaves into powder, after which the powdered plant leaves were stored in a polythene bag and labelled appropriately in preparation for subsequent steps. A small quantity of the powder from Hunteria leaves was kept in a Ziploc bag, labelled, and put away for reference and documentation purposes.

### Methanol Extraction

Manske’s method (Manske and Holmes, 2014) was adopted for extraction. A 500g bag of *Hunteria umbellata* leaves powder was emptied and transferred into a glass jar, and a sufficient volume of methanol was added to soak the plant in. The sample was macerated for 72 hours, after which the leaves were filtered using a Whatman filter to obtain the first methanol filtrate. The filtrate obtained from the leaves was concentrated using a rotary evaporator. The concentrate obtained was kept in a beaker and covered using foil paper. About 5g of the pure methanol extract was kept for reference purposes. The recovered methanol from each concentration was used to resoak the leaves again, and more methanol was added to top off the plant material for the second maceration. The methanol filtrate was subsequently obtained after 24 hours, concentrated, and resoaked till the plant content was exhaustively extracted. All the methanol extract obtained from each maceration was pooled, and one-quarter of this extract was transferred into a sample bottle which was sent for analysis, while the remaining three-quarters proceeded to the alkaloid extraction step.

### Alkaloid Extraction

The methanol extract obtained in the previous step was dissolved in water and repeatedly titrated with 2M sulfuric acid to acidify the solution to pH 2 (tested using a pH meter). Steam distillation was done next on the acidified solution to remove all traces of the methanol, then the acidified Hunteria solution was kept in the refrigerator for 5 days, after which it was taken out and filtered. The filtrate was agitated with 100% chloroform to separate the alkaloidal salts, which are freely soluble in water but not in organic solvents like chloroform. An aqueous and organic phase layer was formed from the previous step. The aqueous phase containing the alkaloidal salt was separated and carefully basified with ammonia to liberate free alkaloids; hence, an aqueous and organic phase was formed. The organic phase containing the free alkaloid was separated using a separating funnel and evaporated to dryness using a rotary evaporator to obtain the crude alkaloids. The crude alkaloid of Hunteria was pooled into a screw-capped sample bottle for analysis.

### LC-MS/MS Analysis

Samples were dissolved in methanol and analysed using an Agilent LC-MS system composed of an Agilent 1260 Infinity HPLC coupled to an Agilent 6530 ESI-Q-TOF-MS operating in positive mode. A Sunfire analytical C18 column (150 × 2.1 mm; i.d. 3.5 μm, Waters) was used, with a flow rate of 250 μL/min and a linear gradient from 5% B (A: H2O + 0.1% formic acid, B: MeOH) to 100% B over 30 min. ESI conditions were set with the capillary temperature at 320 °C, source voltage at 3.5 kV, and a sheath gas flow rate of 10 L/min. The divert valve was set to waste for the first 3 min. There were four scan events: positive MS, a window from m/z 100−1200, and then three data-dependent MS/MS scans of the first, second, and third most intense ions from the first scan event. MS/MS settings were: three fixed collision energies (30, 50, and 70 eV), a default charge of 1, a minimum intensity of 5000 counts, and an isolation width of m/z 2. Purine C5H4N4 [M + H]+ ion (m/z 121.050873) and hexakis(1H,1H,3H-tetrafluoropropoxy)-phosphazene C18H18F24N3O6P3 [M + H]+ ion (m/z 922.009) were used as internal lock masses. Full scans were acquired at a resolution of 11000 (at m/z 922). A permanent MS/MS exclusion list criterion was set to prevent oversampling of the internal calibrant (Ramos *et al*., 2019).

### Computational Molecular Networking

The Global Natural Product Social Molecular Networking (GNPS) website was used as the computational molecular networking tool. The mass spectrometry file data was converted into an mzXML, which is the accepted format by the tool. The GNPS webpage was accessed on a computer system, and account details were entered. From the main GNPS page, the “create molecular network” tab was clicked on, which led to the workflow input page to start networking. The mzXML file was uploaded using the provided tabs, and the following parameters were entered: for Molecular Networking, a precursor ion mass tolerance of 0.02 Da, fragment ion mass tolerance of 0.02 Da, minimum pairs cosine of 0.6, minimum cluster size of 1, score threshold of 0.6, and no filtering of peaks in a 50 Da window; for Spectral Library Search, a precursor ion mass tolerance of 0.02 Da, fragment ion mass tolerance of 0.02 Da, minimum matched peaks of 6, score threshold of 0.6, top hits per spectrum set to 3, and no filtering of peaks in a 50 Da window.

The workflow was then submitted, and the status page displayed, which showed the status of the molecular networking job. On completion, DONE was displayed on the job status page. The result was downloaded from the “View Unique Library Compounds” window. The software Cytoscape was used to view the clusters.

## RESULTS

The data from the obtained chromatogram and the exported CSV file of isolated compounds were extracted to characterise the active constituents of *Hunteria umbellata* (HU) leaves. Eighteen compounds were identified and characterised from the alkaloid and methanol extract.

### Percentage Yield of Extract

The percentage yield of the crude methanol and alkaloid extract can be calculated using the formula shown below.

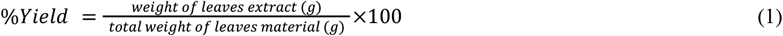

The percentage yield of the crude methanol extract is calculated as:

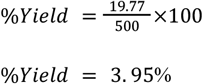

The percentage yield of the alkaloid extract is calculated as:

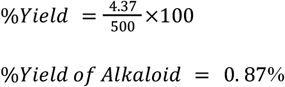

### Isolated Compounds from Methanolic Extract of HU

The LC-MS/MS chromatogram of the methanolic extract of *Hunteria umbellata* (Figure 1) revealed multiple distinct peaks, indicating the presence of several bioactive secondary metabolites. Prominent peaks were observed at retention times of approximately 3.0 and 13.0 minutes, corresponding to high-intensity compounds within the extract. The identified compounds and their analytical characterization data are presented in Table 1.

**Table 1.**
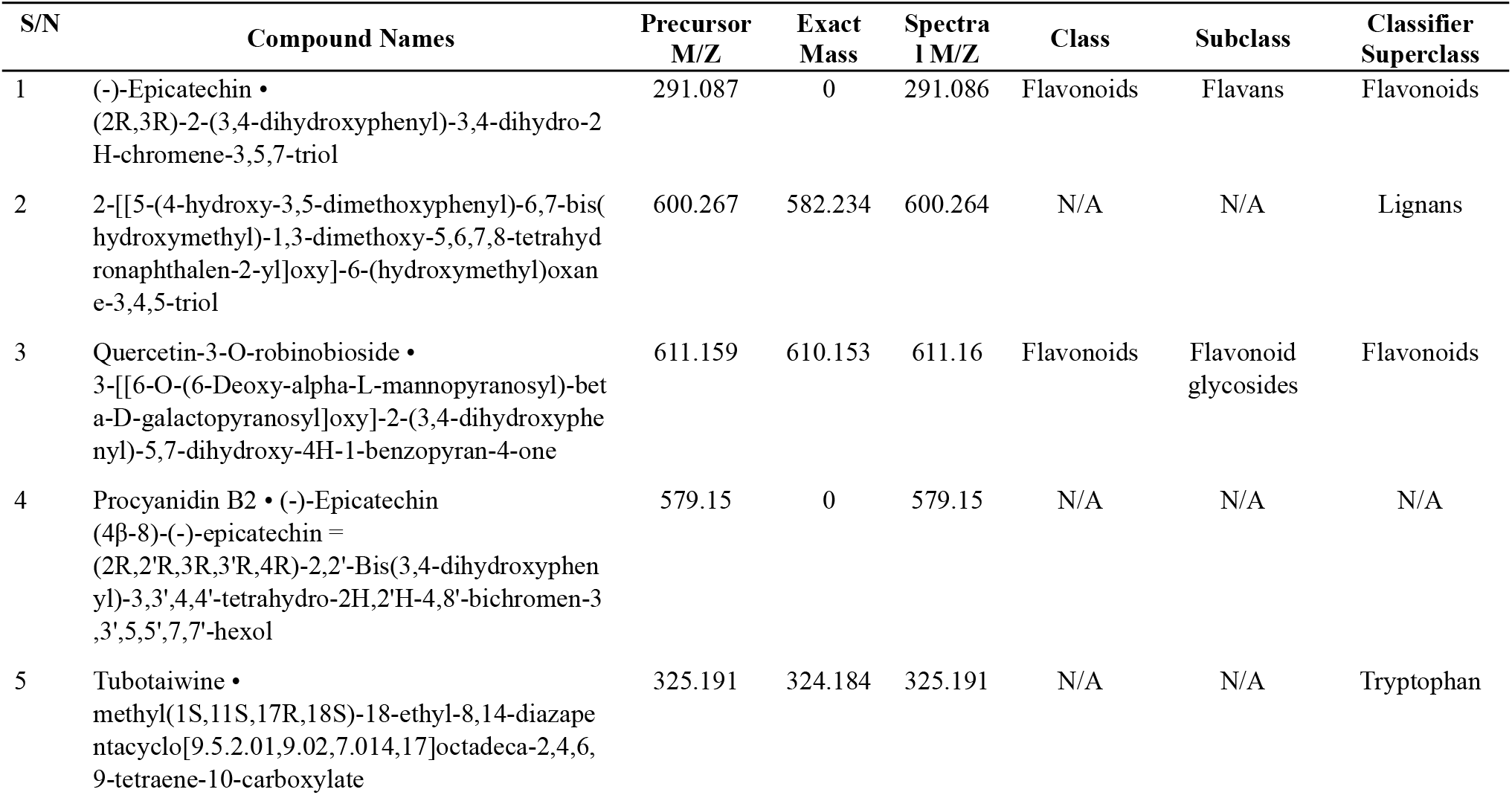
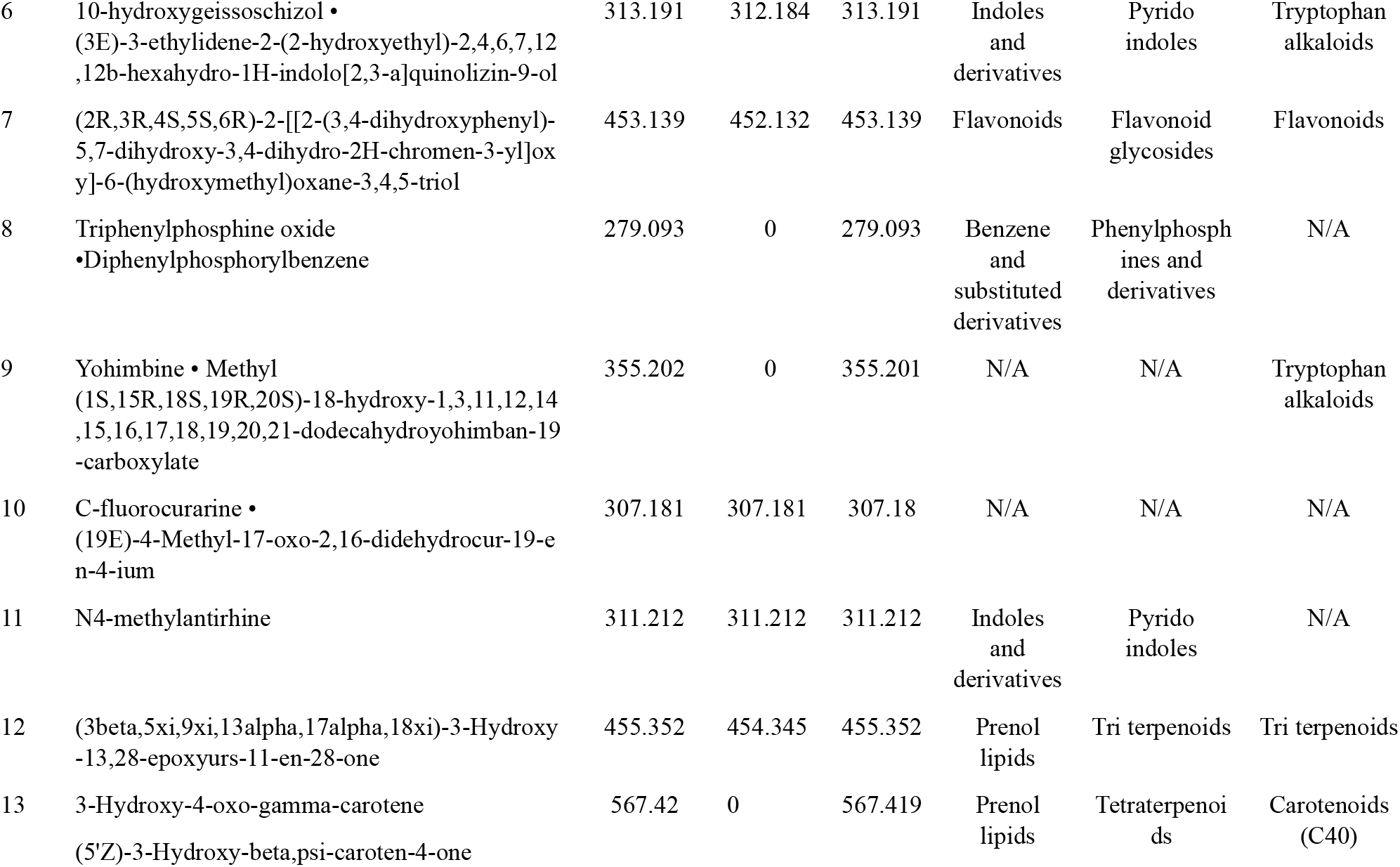
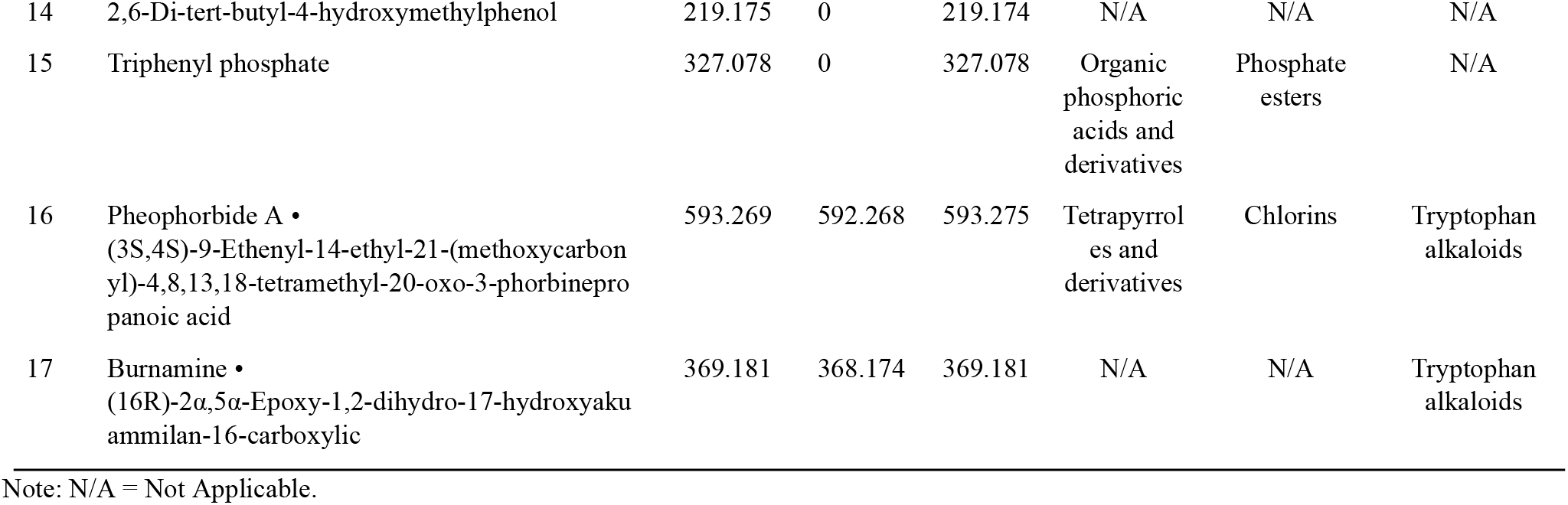
Table of Isolated Compounds from the Methanolic Extract of HU.

**Figure 1.**
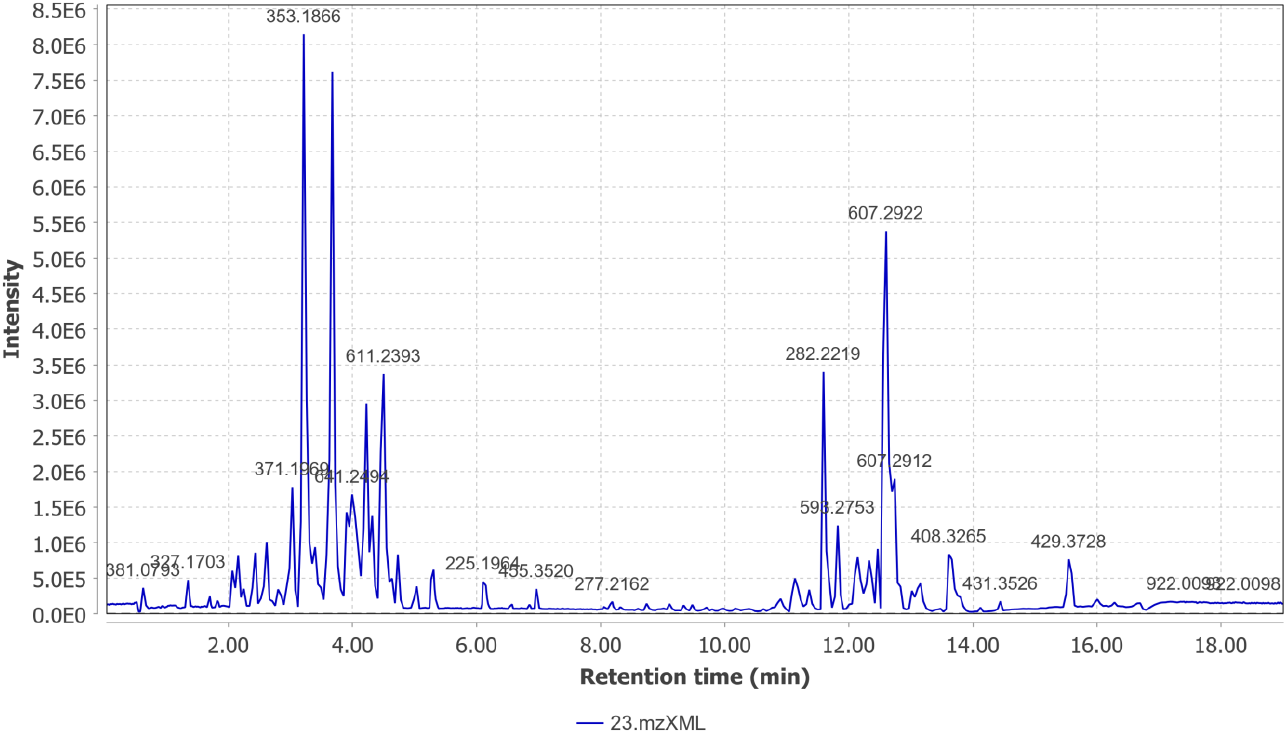
Chromatogram of the Spectrum Peaks of the Isolated Compounds in the Methanolic Extract of HU.

Analysis of the chromatographic data and exported compound tables identified a total of seventeen metabolites (Table 1), belonging predominantly to the flavonoid, alkaloid, lignan, lipid, and carotenoid classes.

Among these, (−)-Epicatechin and Quercetin-3-O-robinobioside were identified as major flavonoids, while Tubotaiwine, Yohimbine, and 10-hydroxygeissoschizol represented key alkaloid constituents. These compounds are known for diverse pharmacological activities, including antioxidant, anti-inflammatory, and antimicrobial effects. The presence of both flavonoid and indole alkaloid derivatives suggests that the methanolic extract of *H. umbellata* contains a complex mixture of bioactive molecules that may collectively contribute to its traditional medicinal uses. The representative structures of selected compounds are shown in Figure 2.

**Figure 2.**
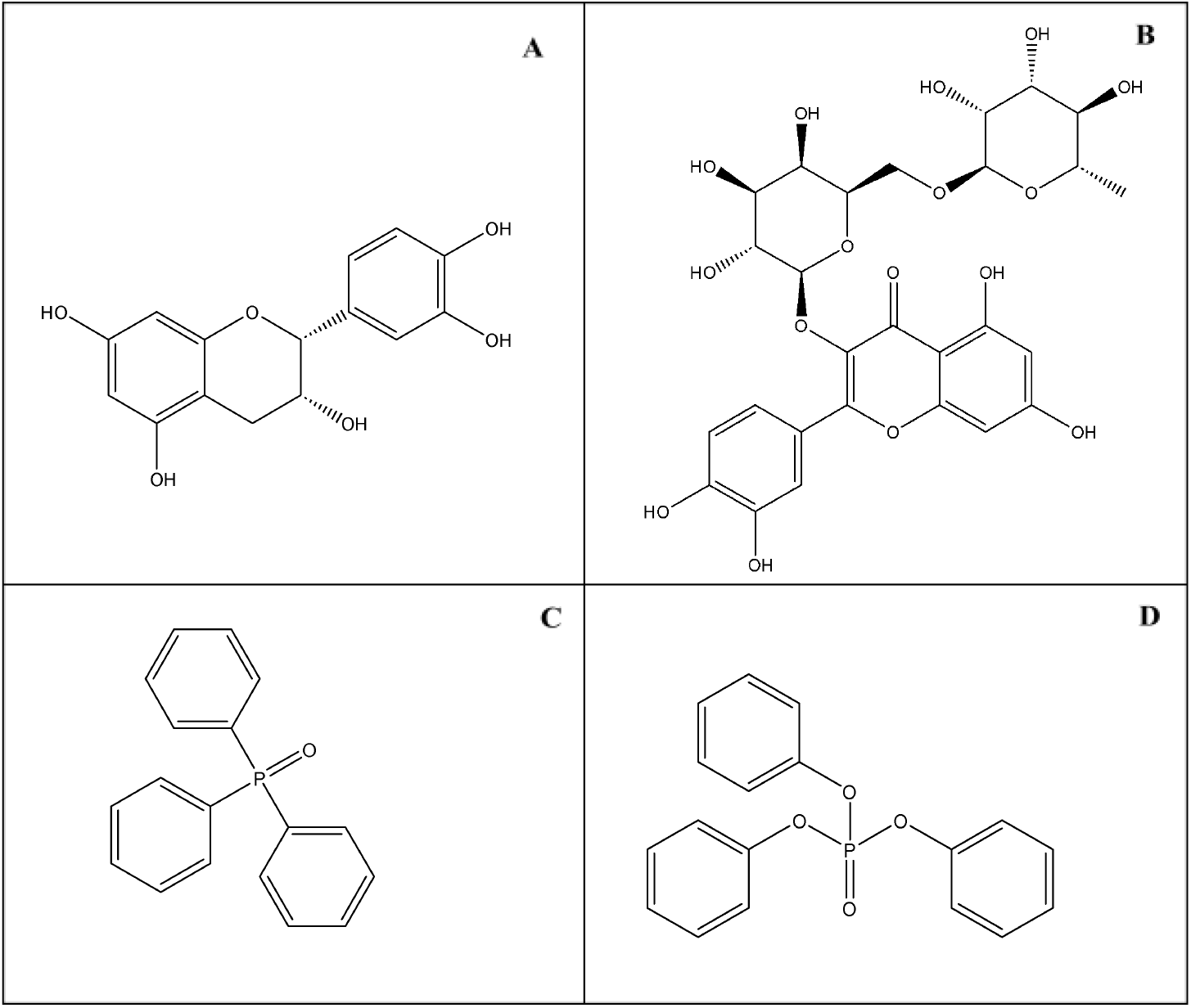
Structures of Some Isolated Compounds Present in the Methanolic Extract. A) (−)-Epicatechin, B) Quercetin-3-O-robinobioside, C) Triphenylphosphine oxide and D) Triphenyl phosphate

### Isolated Compounds from Alkaloid Extract of HU

The LC-MS/MS chromatogram of the alkaloid extract of *Hunteria umbellata* (Figure 3) exhibited several well-defined peaks, indicating the presence of distinct alkaloid and lipid-derived compounds. Prominent peaks were observed between retention times of 11.0 and 13.0 minutes, corresponding to high-intensity molecular ions. The comprehensive analytical data are presented in Table 2, while the corresponding chemical structures are shown in Figure 4 below.

**Table 2.**
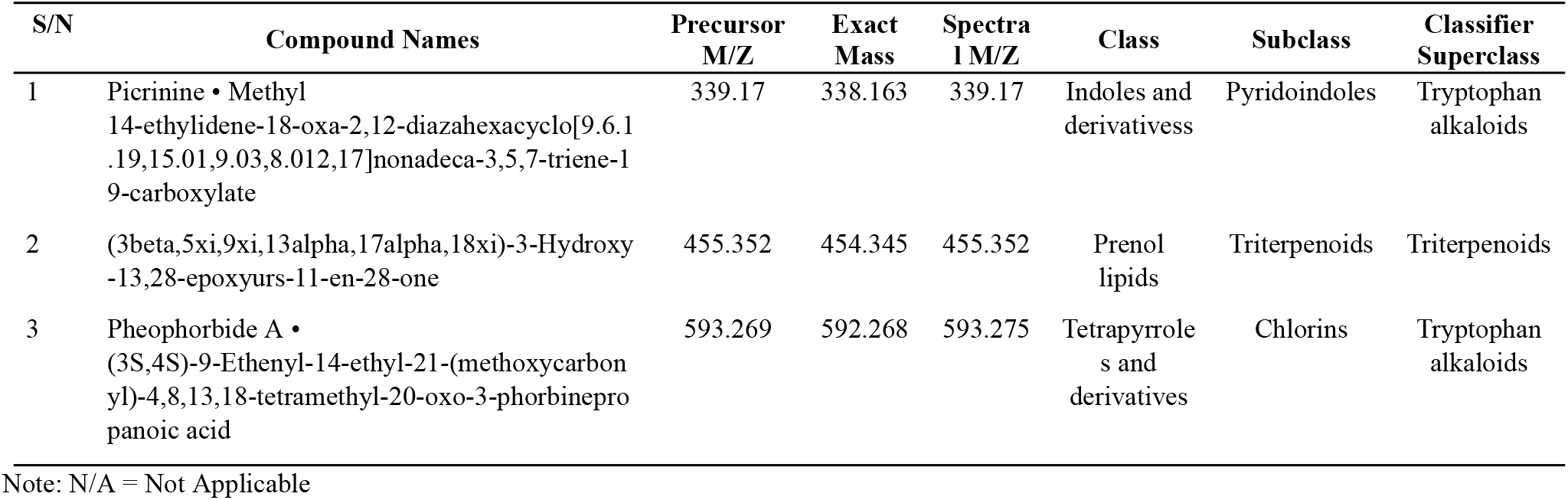
Table of Isolated Compounds from the Alkaloid Extract of HU.

**Figure 3.**
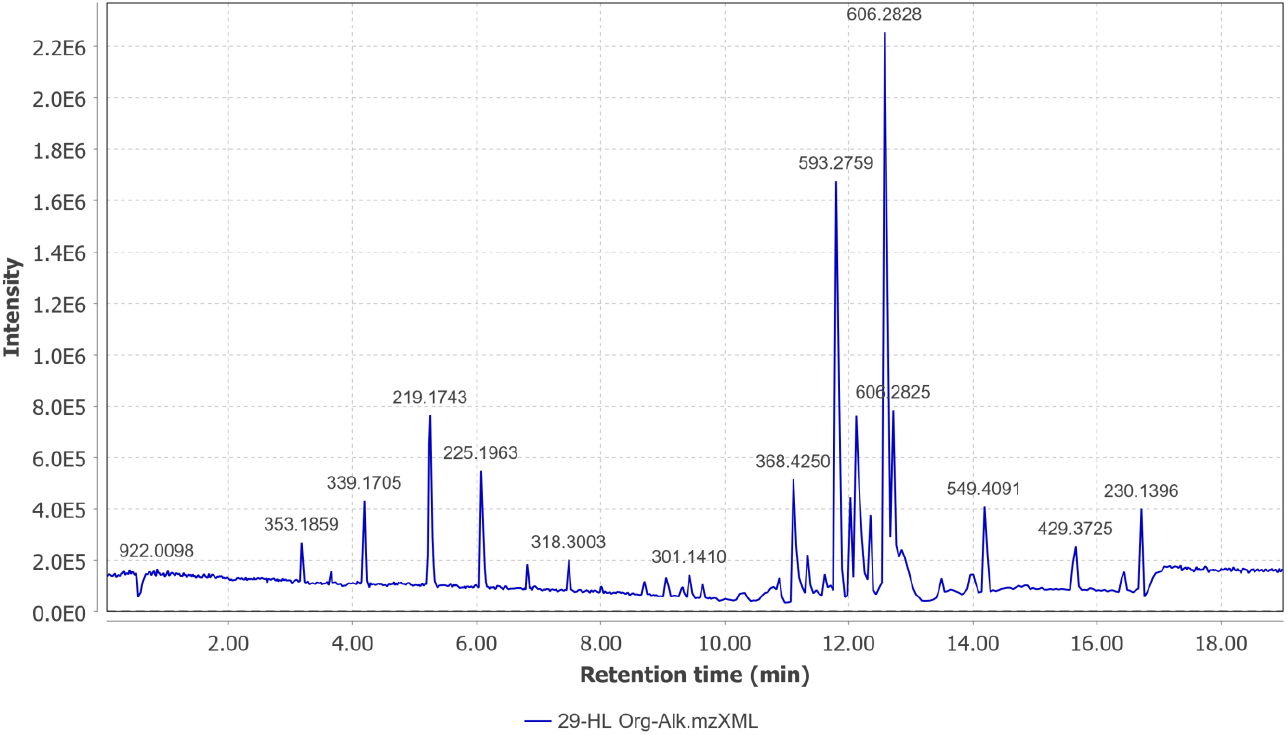
Chromatogram of the Spectrum Peaks of the Isolated Compounds in the Alkaloid Extract of HU.

**Figure 4.**
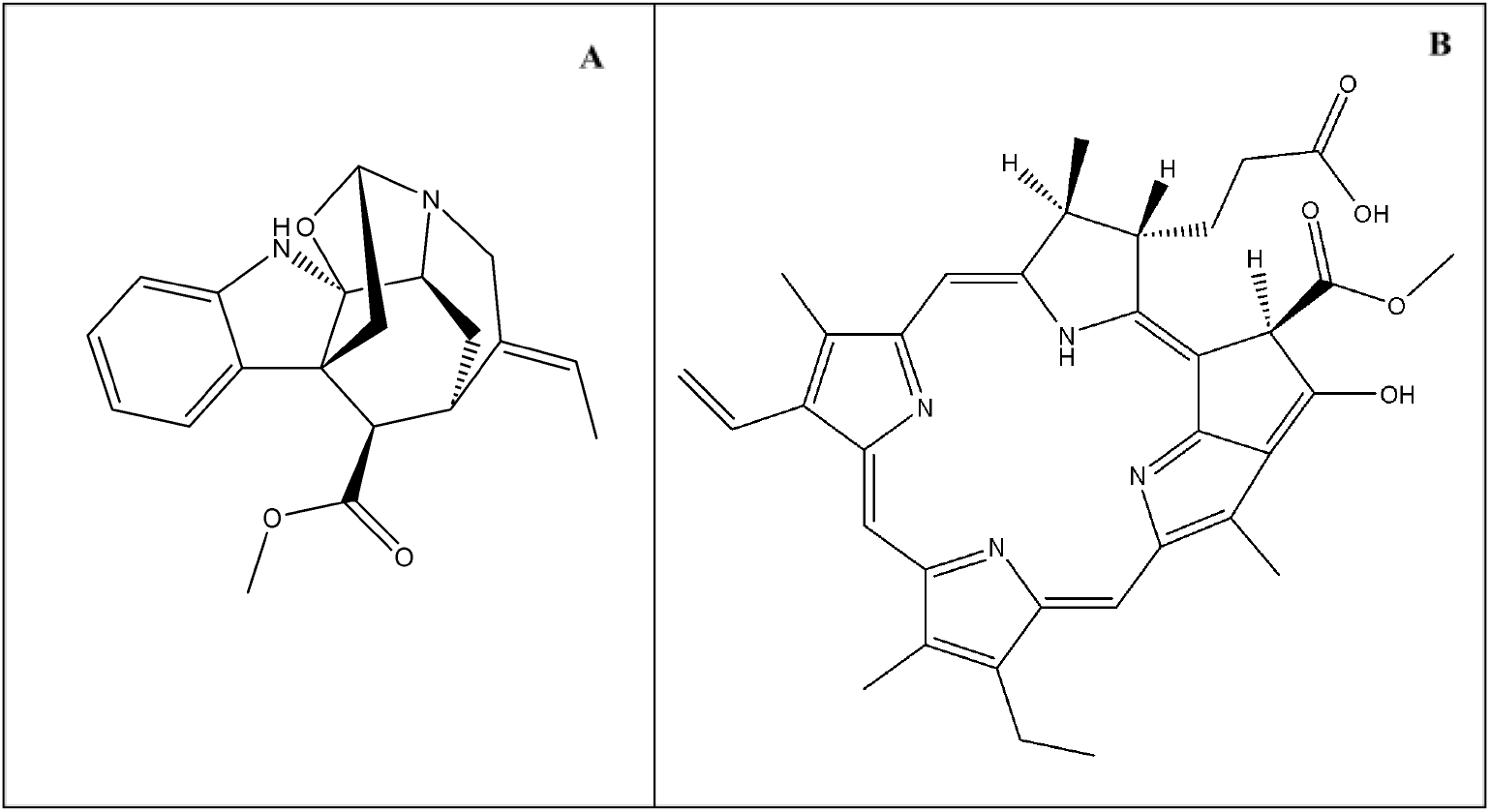
Structures of Some Isolated Compounds Present in the Alkaloid Extract. A) Picrinine, B) Pheophorbide A

Analysis of the chromatographic spectra and GNPS output confirmed the identification of three major compounds. Picrinine, a monoterpenoid indole alkaloid, and Pheophorbide A, a chlorin derivative, were among the most abundant metabolites detected, reflecting the strong representation of tryptophan-derived alkaloids in the extract. The presence of the triterpenoid compound, (3beta,5xi,9xi,13alpha,17alpha,18xi)-3-Hydroxy-13,28-epoxyurs-11-en-28-one suggests that the alkaloidal fraction also contained lipophilic metabolites, possibly due to co-extraction during fractionation. The representative molecular structures of these compounds are illustrated in Figure 4.

## DISCUSSION

Omics technologies coupled with computational tools are revolutionising herbal medicine research as they accelerate the analysis and interpretation of large molecular datasets (Lalit *et al*., 2010; Beale *et al*., 2018). These tools are essential in the development of novel therapeutic leads by implementing algorithms to screen large biomolecule data in silico and effectively predict the efficacy, safety, and pharmacological activity of compounds (Oladipupo *et al*., 2024). Platforms like the Global Natural Products Social Molecular Networking (GNPS), Traditional Chinese Medicine Systems Pharmacology Database and Analysis Platform (TCMSP) are crucial for organising molecular data, promoting data sharing, and driving discoveries. The integration of omics techniques and computational molecular networking for this study allowed us to map out a more refined phytochemical profile of *Hunteria umbellata* beyond what conventional phytochemical screens could achieve. Prior phytochemical surveys of *H. umbellata* typically report the presence of alkaloids, flavonoids, tannins, saponins, cardiac glycosides, and phenolics (Adeneye and Adeyemi, 2009; Igbe *et al*., 2009; Adeneye *et al*., 2011). However, none of those reports mention chlorin-derived molecules like pheophorbide A or deeply annotate networks of low-abundance metabolites.

Our work confirms that *H. umbellata* leaves indeed contain many of the classes previously reported but also adds new dimensions. This suggests that LC-MS/MS with molecular networking offers greater sensitivity to non-volatile, conjugated metabolites that classical screens miss.

The flavonoids isolated from the methanolic extract of HU are; (−)-Epicatechin, Quercetin-3-O-robinobioside, and (2R,3R,4S,5S,6R)-2-[[2-(3,4 dihydroxyphenyl)-5,7-dihydroxy-3,4-dihydro-2H-chromen-3-yl]oxy]-6-(3beta,5xi,9xi,13alpha,17alpha,18xi)-3-Hydroxy-13,28-epoxyurs-11-en-28-one (hydroxymethyl)oxane-3,4,5-triol. The antioxidative effect of catechol-type flavonoids has been extensively studied, and in one such study, (−)-Epicatechin was examined and found to inhibit the accumulation of phosphatidylcholine-hydroperoxides initiated by 2,2′-azobis(2-amidinopropane) hydrochloride (AAPH) in a concentration-dependent manner (Terao *et al*., 1994). The antioxidant and radical-scavenging properties of catechol-type flavonoids are well documented (Terao *et al*., 1994), and these compounds may underpin some of the plant’s ethnopharmacological uses (e.g., anti-inflammatory, analgesic). Also, because of the link between the production of reactive oxygen species (ROS) and inflammation, the antioxidative action of (−)-Epicatechin may also validate the use of HU as an analgesic and anti-inflammatory agent (Adeneye *et al*., 2011).

Recent findings by Álvarez-Cilleros *et al*. (2023) demonstrated that (−)-Epicatechin reduces ROS and inflammatory markers via NOX-4 and p38 signaling pathways, providing a specific antioxidant mechanism. Furthermore, recent comprehensive reviews by Qu *et al*. (2021) and Dash *et al*. (2024) have extensively documented (−)-Epicatechin’s physiological functions, including its insulin-sensitizing effects and mitochondrial protective properties (Qu *et al*., 2021; Dash *et al*., 2024). These mechanisms collectively improve insulin sensitivity and highlight the compound’s potential antidiabetic activity. (Álvarez-Cilleros *et al*., 2017; Mechchate *et al*., 2021). These findings align with ethnobotanical reports of HU in diabetes management, suggesting that this flavonoid contributes significantly to the antidiabetic properties of the plant through multiple mechanistic pathways.

The alkaloids identified in this study include; Tubotaiwine, 10-hydroxygeissoschizol, Yohimbine, Pheophorbide A, Burnamine, Picrinine and (3beta,5xi,9xi,13alpha,17alpha,18xi)-3-Hydroxy-13,28-epoxyurs-11-en-28-one. Tubotaiwine, a tryptophan alkaloid, has been studied and found to possess antimicrobial activity among other activities, which may explain why the plant is traditionally used in the treatment of infection (Barati and Chahardehi, 2023). Recent antimicrobial screening of HU seed extracts by Salisu *et al*. (2024) has shown significant bactericidal activity against *Escherichia coli, Staphylococcus aureus*, and *Streptococcus species*, with measured minimum inhibitory concentrations supporting its traditional antimicrobial applications (Salisu *et al*., 2024). Another biological study found tubotaiwine to be an opioid receptor ligand, which might suggest some psychoactive or analgesic activity, corroborating its local use for pain management (Stafford *et al*., 2009).

The identification of Yohimbine which is a well-characterized alpha-2 adrenergic antagonist in the methanolic extract provides molecular support for the traditional use of HU in treating male sexual dysfunction. This finding correlates with recent pharmacological studies showing the aphrodisiac effects of HU in streptozotocin-induced diabetic rats (Ogunlana *et al*., 2021). A study by Oboh *et al*. (2019) found that administration of *Hunteria umbellata* seed extract in male rats increased nitric oxide and antioxidant levels, as well as improved sexual behaviour parameters, supporting its traditional use as an aphrodisiac. Recent receptor binding studies by Proudman *et al*. (2022) confirmed Yohimbine as a high-affinity, non-selective α2 antagonist with precise quantitative binding data for human α2A/α2B/α2C subtypes(Proudman *et al*., 2022). However, a review by Nowacka *et al*. (2024) emphasize important safety considerations and potential herb-drug interactions that warrant careful evaluation in therapeutic applications (Nowacka *et al*., 2024).

The identification of pheophorbide A is particularly interesting because it lies outside the typical spectrum of secondary metabolites documented for *H. umbellata*. Pheophorbide A is well-established in photodynamic therapy (PDT) research as a potent photosensitizer capable of inducing ROS-mediated cytotoxicity in cancer cell lines (Chen *et al*., 2009; Bui-Xuan *et al*., 2010). Also, Gheewala *et al*. (2023) demonstrated that Pheophorbide A-mediated photodynamic therapy induces ROS generation, ER stress, and apoptosis in cancer cell lines (Gheewala *et al*., 2023). These literatures validate that pheophorbide A is not merely a pigment derivative but a biologically active molecule with therapeutic potential. Although we cannot yet ascribe a cancer-therapeutic role to its presence in HU, its detection is a new report for *H. umbellata* and highlights avenues for future investigation.

Burnamine has been investigated for hypoglycemic activity, but no significant activity has been recorded to validate the use of the HU plant for glycemic control, unlike Akuammicine, a structurally similar monoterpenoid indole alkaloid (Erharuyi *et al*., 2014). While structurally similar alkaloids in other Apocynaceae sometimes show insulin-modulating activity, there is currently no strong published evidence that burnamine itself exerts significant glucose-lowering effects when isolated.

A study by Onygeme-Okerenta *et al*. (2023) showed that picrinine possesses anti-inflammatory, antitussive and expectorant activity (Onygeme-Okerenta *et al*., 2023). Hence, the presence of picrinine in the alkaloid extract aligns with the traditional use of HU in treating respiratory conditions.

Procyanidine B2 is a B-type proanthocyanidin that was also isolated. A laboratory study suggests that Procyanidin B2 plays an important role in hair cycle progression and may be useful in promoting hair growth (Kamimura and Takahashi, 2002).

There have been suggestions that C-fluorocurarine possesses a hypotensive effect and might be useful in treating hypertension, but these claims are yet to be validated by strong scientific evidence. 2,6-Di-tert-butyl-4-hydroxymethylphenol is another phenolic compound isolated from HU leaves and reported to possess antioxidant action but will require more research to confirm that and its other activities. The other compounds isolated are relatively novel moieties, with limited information currently available. Further studies are required to determine their medicinal relevance or potential cytotoxicity.

In closing, while our study significantly advances the chemical understanding of *Hunteria umbellata*, cautious interpretation and further validation are vital.

## CONCLUSIONS

This study provides a comprehensive LC-MS/MS-based metabolomic characterisation of the methanolic and alkaloidal extracts of *Hunteria umbellata* leaves, revealing both known and previously unreported bioactive compounds. Eighteen compounds were identified, spanning major phytochemical classes such as alkaloids, flavonoids, triterpenoids, and chlorins. Notably, the identification of pheophorbide A represents the first report of this chlorin derivative in *H. umbellata* and suggests unexplored therapeutic potential, particularly in photodynamic cancer therapy. The detection of (−)-epicatechin and quercetin derivatives also provides a basis for the plant’s antidiabetic and antioxidant traditional uses, while the presence of yohimbine corroborates its aphrodisiac applications. Compared to previously studied aqueous and ethanolic extracts, which primarily revealed common alkaloids and flavonoids, the present findings demonstrate the superior sensitivity of LC-MS/MS and molecular networking approaches in unveiling deeper chemical complexity. Overall, this work advances the phytochemical understanding of *Hunteria umbellata*, highlights key metabolites as promising leads for future drug discovery efforts and validates the integration of traditional knowledge with modern analytical approaches in natural product research.

## Data Availability

Data that supports the reported results of this research is available from the corresponding author.

